# Open-source benchmarking of dairy and dairy-free products

**DOI:** 10.64898/2026.08.14.744929

**Authors:** Aeneas O. Koosis, Caroline Cotto, Ellen Kuhl

## Abstract

Dairy-free products offer opportunities to reduce the environmental impact of food. However, they must reproduce the diverse sensory and functional properties of conventional dairy, from foam stabilization in a latte to acid-gel structure in yogurt and melt– stretch behavior in mozzarella. Yet, large-scale, open-source sensory benchmarks across dairy product categories remain scarce. Here we show that the sensory gap between dairy and dairy-free products varies substantially across ten categories: barista milk, butter, cheddar, cream cheese, creamer, ice cream, fluid milk, mozzarella, sour cream, and Greek-style yogurt. We analyze the NEC-TAR Taste of the Industry 2026 dataset, which includes 1,437 respondents, 3,220 overall-liking ratings, and 79,027 recorded survey responses, including 35,762 product-linked responses. Leading dairy-free beverages, barista milk (+0.17), fluid milk (+0.02), and creamer (−0.06), closely approach their dairy benchmarks in overall liking on a seven-point scale. In contrast, mozzarella shows the largest deficit (−1.46), followed by Greek-style yogurt (−0.68) and butter (−0.54). All three deficits remain statistically significant after correction for multiple comparisons. Across categories, descriptive penalty analysis identifies off-flavor (1.85 points), stale or cardboard notes (1.81 points), and artificial or chemical notes (1.80 points) as the sensory attributes associated with the largest reductions in overall liking. Together, these results establish the first open sensory benchmark for dairy and dairy-free products, identify beverage categories that approach conventional dairy in overall liking, and reveal the sensory attributes and product categories with the greatest opportunities for future innovation.

## 1. Introduction

Food production contributes substantially to greenhouse-gas emissions, land conversion, freshwater use, and biodiversity loss [6, 18]. Dietary change could reduce these environmental pressures alongside improvements in food production [22, 27, 5]. Plant-based milks generally use less land and produce fewer greenhouse-gas emissions than cow’s milk, although estimates vary by crop, geography, allocation method, and functional unit [9]. These environmental advantages, however, do not ensure that consumers will replace dairy in routine meals [12].

Conventional dairy products provide important nutrients, including high-quality protein, calcium, and vitamins, that contribute to normal growth and bone health [16]. National dietary guidelines commonly recommend dairy consumption, although recommendations vary across countries and income groups [3]. At the same time, demand for dairy alternatives continues to grow, driven by lactose intolerance, milk-protein allergies, dietary preferences, and concerns about health, animal welfare, and environmental sustainability [4]. Yet, replacing dairy is not a simple substitution problem: Fluid milk is an oil-in-water emulsion and colloidal protein dispersion; steamed milk is also a transient foam. Yogurt is an acid-induced protein gel, butter is a water-in-oil emulsion structured by a partially crystalline fat network, and ice cream contains ice crystals, air cells, fat globules, and an unfrozen continuous phase. Cheese adds a calcium-associated casein network whose response to heating controls flow, stretch, oil retention, and fracture [15, 19, 29, 14, 13]. Plant proteins, starches, vegetable fats, and hydrocolloids can reproduce portions of these structures, but they do not inherit the same interfacial, gelation, crystallization, or flavor-release behavior.

The commercial record reflects this difference in technical difficulty. Plant-based milk generated about one third of U.S. plant-based retail dollars and 13% of total milk dollar sales in 2025 [7]. Consumer studies identify smoothness, creaminess, and sweetness as positive drivers of liking in plant-based milks, with barista-style products among the better-performing formats [8, 1]. In coffee, performance depends not only on beverage viscosity but also on protein adsorption at the air–water interface, liquid drainage between bubbles, heat stability, and the shear history imposed during steaming. Model cappuccino systems show that hydrocolloid-controlled drainage can markedly alter plant-protein foam stability [28]. Structured products pose a different problem. Most plant proteins form heat-set aggregates or brittle networks rather than the thermoreversible, extensible matrix required for mozzarella; zein-based systems can improve stretch, but melt and cohesion remain formulation-sensitive [14, 21]. Fermented alternatives must also generate a continuous gel while controlling syneresis and raw-material-derived volatiles [29, 26].

Consumers of dairy alternatives cite taste alongside health and environmental considerations, and sensory work repeatedly shows that acceptance depends on the product and use context rather than on a general attitude toward plant-based foods [8, 1]. This makes direct tasting against a familiar dairy reference especially informative. It also raises a measurement problem: a small, non-significant difference in a conventional superiority test is not evidence that two products are sensorially equivalent. Equivalence requires a justified sensory margin and a test designed around that margin. Large proprietary datasets could support such analyses, but they are seldom available for independent scrutiny.

Here we analyze the NECTAR Taste of the Industry 2026 dataset [17] and compare dairy benchmarks with selected dairy-free category leaders from blinded consumer tastings across ten product categories. Our study builds on a companion bench-mark of plant-based and animal meats [2]. *We hypothesize that the sensory gap between leading dairy-free products and their dairy counterparts varies systematically across product categories and reflects distinct sensory attributes associated with consumer acceptance*. To test this hypothesis, we analyze paired differences in consumer ratings, identify sensory attributes associated with lower liking, and compare sensory performance across dairy categories and serving contexts. Together, these analyses establish a quantitative sensory benchmark that helps researchers and product developers identify opportunities for improvement and guide the design of future dairy-free products.

## 2. Materials and methods

### 2.1. Study design

We analyze the NECTAR Taste of the Industry 2026 dataset [17], where NECTAR designed the source study, Palate Insights collected the data, and we performed a secondary analysis. Palate Insights conducted blinded in-person sensory tests at restaurant partner locations in San Francisco, CA, and New York City, NY, between September 2025 and November 2025. Participants evaluated products one at a time under blinded and randomized conditions and recorded their sensory experience in a survey. Products were prepared according to manufacturer instructions. More complex dishes underwent trial preparations before testing to confirm visual standards with manufacturers.

The full study tested 98 unique commercially available dairyfree products and ten conventional dairy benchmarks across barista milk, butter, cheddar, cream cheese, creamer, ice cream, fluid milk, mozzarella, sour cream, and Greek-style yogurt (Fig. 1). Four exploratory products that combined plant and conventional dairy ingredients were also tested in the source study but were excluded here. NECTAR selected the *dairy benchmark* as the highest-volume product in the retail multioutlet channel and the *dairy-free leader* post hoc as the product with the highest overall liking score [17]. The final sample contained 1,437 respondents, 3,220 overall-liking ratings, and 79,027 recorded survey responses for the ten dairy benchmarks and the ten selected dairy-free leaders.

**Figure 1:**
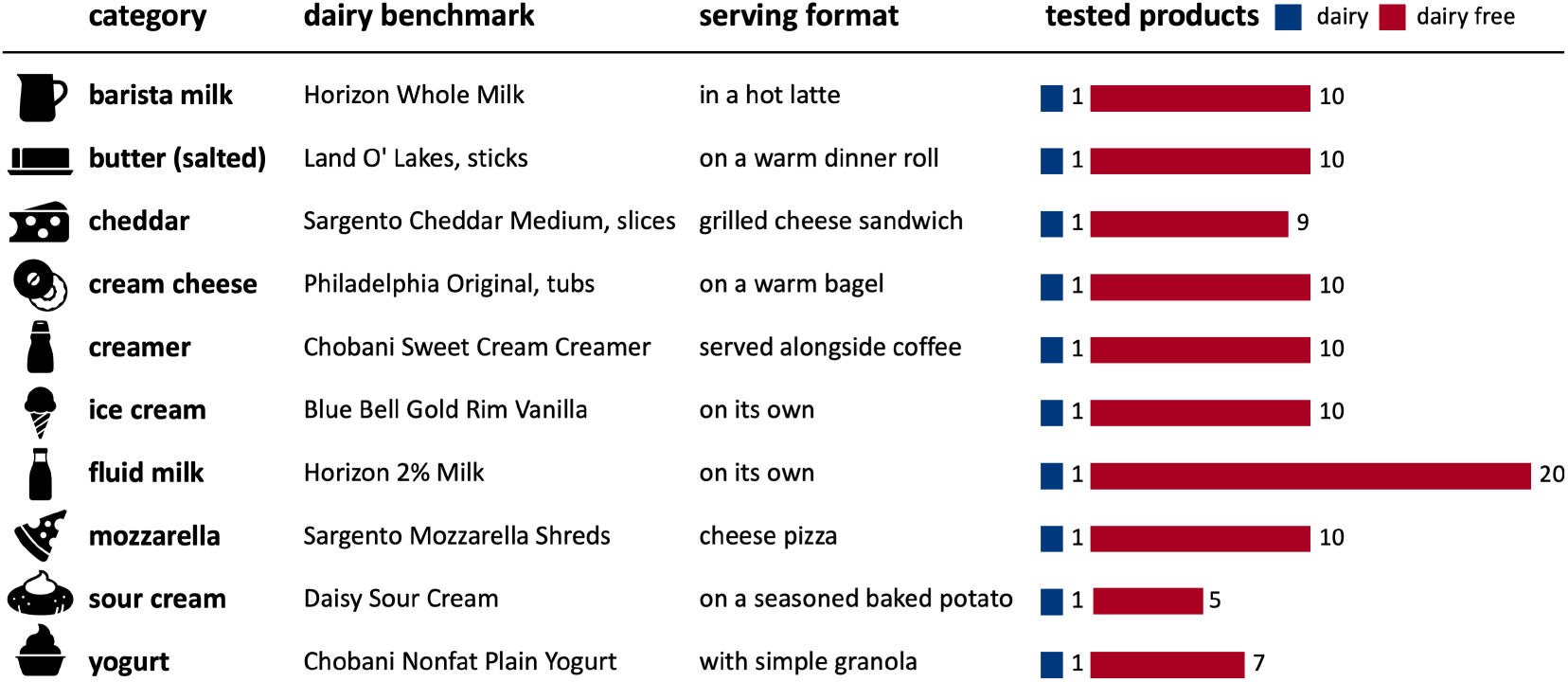
Product categories and serving formats. The study evaluated ten dairy categories, each represented by a highest-volume dairy benchmark and a set of commercially available dairy-free products (108 unique products: 98 dairy-free and 10 benchmarks; balanced-dairy products excluded). Bars show the number of dairy-free products (red) and dairy benchmarks (dark blue) evaluated per category. Dairy benchmarks were selected by retail sales volume in the multi-outlet (MULO) channel to represent the typical dairy product, and each product was served in a complete but simplified, category-specific dish.

### 2.2. Product categories and serving formats

Figure 1 summarizes the dairy categories, benchmark products, and serving contexts reported by NECTAR [17].

### 2.3. Study population

The final sample contains 1,437 respondents, including omnivores, flexitarians, vegetarians, and pescatarians [17]. These respondents provided 2,183 benchmark ratings and 1,037 leader ratings. The paired analysis includes 1,037 comparisons from 725 respondents, and 170 of these respondents contributed pairs in more than one category. The demographic data include gender, age, dietary preference, education, ethnicity, household composition, product-category consumption frequency, current dairy versus dairy-free consumption, shopping priorities, and attitudes toward taste, price, health, environment, familiarity, and animal welfare.

### 2.4. Sensory and consumer measures

Participants rated overall liking, flavor, texture and mouthfeel, appearance, and similarity to dairy on seven-point scales under blinded conditions. After reviewing the nutrition facts panel and ingredient list, they rated purchase intent on a sevenpoint scale. They also completed check-all-that-apply (CATA) questions for appearance, flavor, and texture/mouthfeel attributes and free-response questions about likes, dislikes, and off-flavors or aftertastes. The survey further included pricesensitivity questions, naming questions, and sources that influenced opinions about dairy-free products.

### 2.5. Statistical analysis

The primary analysis compares each *dairy benchmark* with the selected *dairy-free leader*. The available data include observations for each category’s benchmark and selected leader, but not for the remaining dairy-free products. We therefore could not reproduce leader selection, estimate selection optimism, or characterize category-average dairy-free performance independently. Products that combine conventional and plant-derived dairy ingredients were excluded from all analyses.

For each category and modality, we calculated paired means and mean leader-minus-benchmark differences with 95% *t*-based confidence intervals among the *n* = 96–114 participants who rated both products. The study aimed for *n* = 100 participants per product, so pair counts varied around this target. Paired *t* tests were the primary tests of mean differences; two-sided Wilcoxon signed-rank tests with zero differences omitted were sensitivity analyses and produced the same Holm-adjusted overall-liking decisions. The ten overall-liking comparisons constituted the primary superiority family, for which we controlled the family-wise error rate with the Holm procedure. We also conducted exploratory category-specific two one-sided tests (TOST) with a chosen equivalence margin of ±0.5 points on the seven-point scale; equivalence required the 90% confidence interval for the paired mean difference to lie wholly within the margin. These category-specific TOST results were not adjusted for multiplicity. As sensitivity checks, we repeated the TOST with margins from ±0.25 to ±1.0 points and applied Holm and Bonferroni corrections across the ten cat-egories. For the 40 exploratory component-rating comparisons, we controlled the false-discovery rate using the Benjamini– Hochberg procedure. We report Cohen’s *d*_*z*_ for paired mean differences and shares of ratings in three descriptive bands: 6– 7, 4–5, and 1–3. To examine age, we modeled paired overall-liking differences across the five age groups for 1,037 paired observations from 725 respondents. We fit a linear model with category and age group as factors and respondent-clustered standard errors, and we tested the age effect and the category-by-age interaction with Wald *F* tests. Within each category, we compared age groups with one-way ANOVA and applied the Holm procedure across the ten categories.

For each CATA term, we report prevalence and a descriptive penalty: the mean overall liking among participants who did not select the term minus the mean among those who did. We computed penalties per category for each dairy-free leader among terms selected for at least ten ratings and pooled across leaders for terms with at least 25 mentions. Percentile 95% confidence intervals were obtained from 10,000 bootstrap resamples of the selected and unselected rating groups. The bootstrap did not adjust for category or repeated participation, so the intervals do not account for respondents who rated more than one leader, and the penalties represent associations rather than causal effects of individual attributes. Two stated price thresholds were recorded per category: the price at which the product would be considered a bargain and the price at which it would be considered expensive but still purchasable. Analyses were conducted in Python using de-identified observations.

## 3. Results

### 3.1. Study population

The sample skews female (56%) and young to middle-aged, with 60% between 18 and 35 years of age; 66% identify as omnivores and 25% as flexitarians; 49% hold a bachelor’s degree and a further 23% a master’s or doctoral degree; and 12% report children in the household. This composition reflects a consumer panel recruited among regular category consumers rather than a demographically representative population.

### 3.1. Overall liking and equivalence across dairy categories

Paired overall-liking differences, dairy-free leader minus dairy benchmark, ranged from −1.46 to +0.17 points across categories (Fig. 2, Table 2). Mozzarella had a mean paired difference of −1.46 points (*p* < 0.001; *d*_*z*_ = −0.88), followed by yogurt (−0.68; *p* = 0.002; *d*_*z*_ = −0.31) and butter (−0.54; *p* = 0.002; *d*_*z*_ = −0.31). These three comparisons remained significant after Holm correction across the ten primary tests, and the Wilcoxon sensitivity analysis reproduced every Holm decision. Cream cheese had a mean paired difference of −0.42 points (*p* = 0.020) and did not remain significant after correction. The remaining categories had mean paired differences from −0.28 to +0.17 points and |*d*_*z*_| 0.12.

**Table 1:** Study population. Demographic composition of 1,437 respondents

| Characteristic | % | Characteristic | % |
|---|---|---|---|
| *Gender* |  | *Education* |  |
| Female | 56.1 | Some high school | 0.5 |
| Male | 38.9 | High school | 6.8 |
| Non-binary | 3.5 | Trade school | 1.1 |
| Prefer not to say | 1.5 | Some college | 19.5 |
| *Age* |  | Bachelor’s degree | 49.3 |
| 18–25 | 27.3 | Master’s degree | 18.6 |
| 26–35 | 32.9 | Ph.D. or higher | 4.2 |
| 36–45 | 16.2 | *Diet* |  |
| 46–55 | 10.9 | Omnivore | 66.2 |
| Over 55 | 12.6 | Flexitarian | 24.5 |
| *Children in household* |  | Vegetarian | 5.9 |
| Yes | 11.9 | Pescatarian | 3.3 |
| No | 88.1 |  |  |

**Table 2:**
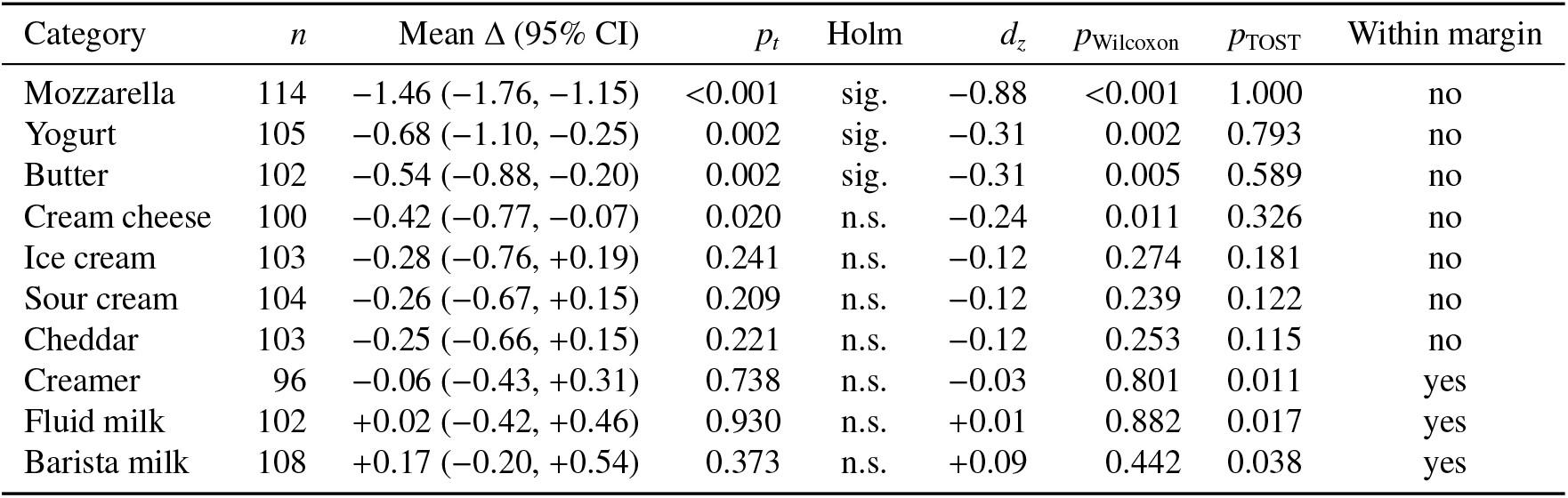
Overall liking: paired benchmark–leader comparisons and exploratory equivalence tests. Mean paired differences are dairy-free leader minus dairy benchmark. The table reports paired *t*-test results, Holm decisions across the ten primary superiority tests, Cohen’s *d*_*z*_, Wilcoxon sensitivity results, and unadjusted category-specific TOST results against a chosen ±0.5-point margin. Mean differences carry 95% confidence intervals; equivalence decisions use 90% confidence intervals, which Table S1 reports with unadjusted, Holm-adjusted, and Bonferroni-adjusted TOST *p* values.

| Category | *n* | Mean Δ (95% CI) | *p<sub>t</sub>* | Holm | *d<sub>z</sub>* | *p*<sub>Wilcoxon</sub> | *p*<sub>TOST</sub> | Within margin |
|---|---|---|---|---|---|---|---|---|
| Mozzarella | 114 | −1.46 (−1.76, −1.15) | <0.001 | sig. | −0.88 | <0.001 | 1.000 | no |
| Yogurt | 105 | −0.68 (−1.10, −0.25) | 0.002 | sig. | −0.31 | 0.002 | 0.793 | no |
| Butter | 102 | −0.54 (−0.88, −0.20) | 0.002 | sig. | −0.31 | 0.005 | 0.589 | no |
| Cream cheese | 100 | −0.42 (−0.77, −0.07) | 0.020 | n.s. | −0.24 | 0.011 | 0.326 | no |
| Ice cream | 103 | −0.28 (−0.76, +0.19) | 0.241 | n.s. | −0.12 | 0.274 | 0.181 | no |
| Sour cream | 104 | −0.26 (−0.67, +0.15) | 0.209 | n.s. | −0.12 | 0.239 | 0.122 | no |
| Cheddar | 103 | −0.25 (−0.66, +0.15) | 0.221 | n.s. | −0.12 | 0.253 | 0.115 | no |
| Creamer | 96 | −0.06 (−0.43, +0.31) | 0.738 | n.s. | −0.03 | 0.801 | 0.011 | yes |
| Fluid milk | 102 | +0.02 (−0.42, +0.46) | 0.930 | n.s. | +0.01 | 0.882 | 0.017 | yes |
| Barista milk | 108 | +0.17 (−0.20, +0.54) | 0.373 | n.s. | +0.09 | 0.442 | 0.038 | yes |

**Figure 2:**
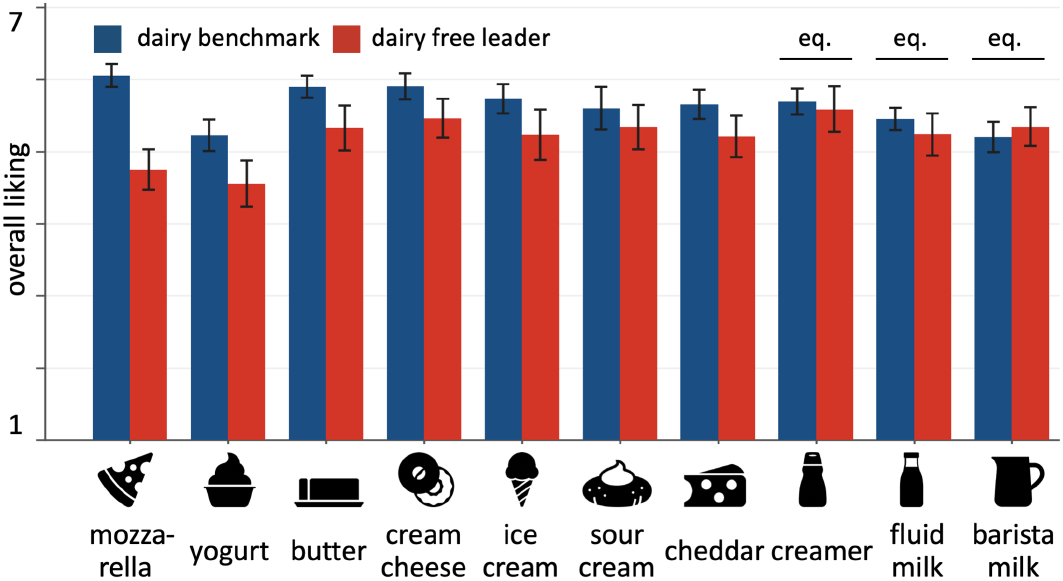
Overall liking by category. Paired-subset mean overall liking (1–7) with 95% confidence intervals for the dairy benchmark (blue) and dairy-free leader (red). TOST annotations identify the three categories whose 90% confidence interval for the paired difference fell within the chosen ±0.5-point margin; eq. denotes unadjusted exploratory equivalence. Categories are ordered by the paired leader-minus-benchmark difference.

The exploratory equivalence tests separated some nonsignificant results, which a superiority test alone cannot do. Us-ing the chosen ±0.5-point margin, creamer (*p*_TOST_ = 0.011), fluid milk (*p*_TOST_ = 0.017), and barista milk (*p*_TOST_ = 0.038) met the category-specific criterion. The margin was not externally validated and these tests were not adjusted for multiplicity, so the results are presented as exploratory evidence of closeness rather than definitive sensory equivalence. The remaining seven categories did not meet the equivalence criterion. None of the three categories retained equivalence after Holm or Bonferroni correction across the ten categories, and the number of categories that met the criterion varied with the margin (Tables S1 and S2). These checks cannot validate the margin or correct for leader selection.

### 3.3. Attribute-level performance

Component-rating differences varied by category (Fig. 3). Of the 40 paired comparisons, 15 remained significant after Benjamini–Hochberg control. Barista milk and creamer had flavor and texture differences within ± 0.10 points and no component difference surviving correction. Mozzarella differed on every component, from −1.22 for appearance to −1.44 for flavor. Fluid milk showed the widest separation between similar-ity and liking: similarity to dairy differed by −1.74 points and appearance by −0.75, while flavor with −0.14 and texture and mouthfeel with −0.28 showed smaller differences.

**Figure 3:**
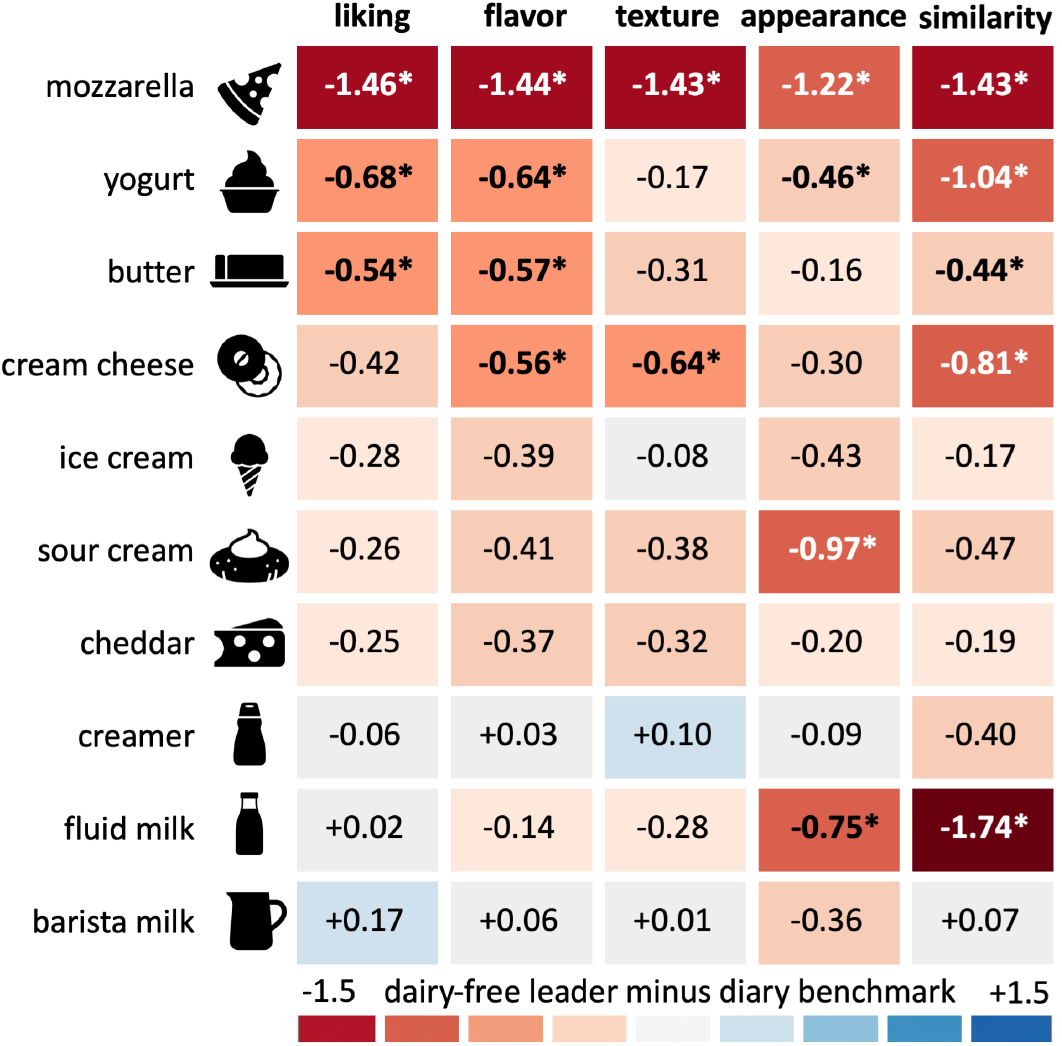
Sensory-gap heatmap. Paired mean difference, dairy-free leader minus dairy benchmark, on 1–7 scale for overall liking, flavor, texture and mouth-feel, appearance, and similarity to dairy. Red shading marks deficits and blue shading marks advantages; shading saturates at |Δ| = 1.5. Asterisks mark component comparisons significant at a Benjamini–Hochberg false discovery rate of 5%; the overall-liking column reproduces Table 2.

### 3.4. Distribution of overall-liking ratings

The descriptive rating bands show where the distributions differ (Fig. 4). Ratings of 6–7 account for 69% of creamerleader evaluations and 59% of sour-cream-leader evaluations, compared with 35% for mozzarella and 30% for yogurt. Ratings of 1–3 account for 24% and 29% of leader evaluations in mozzarella and yogurt.

**Figure 4:**
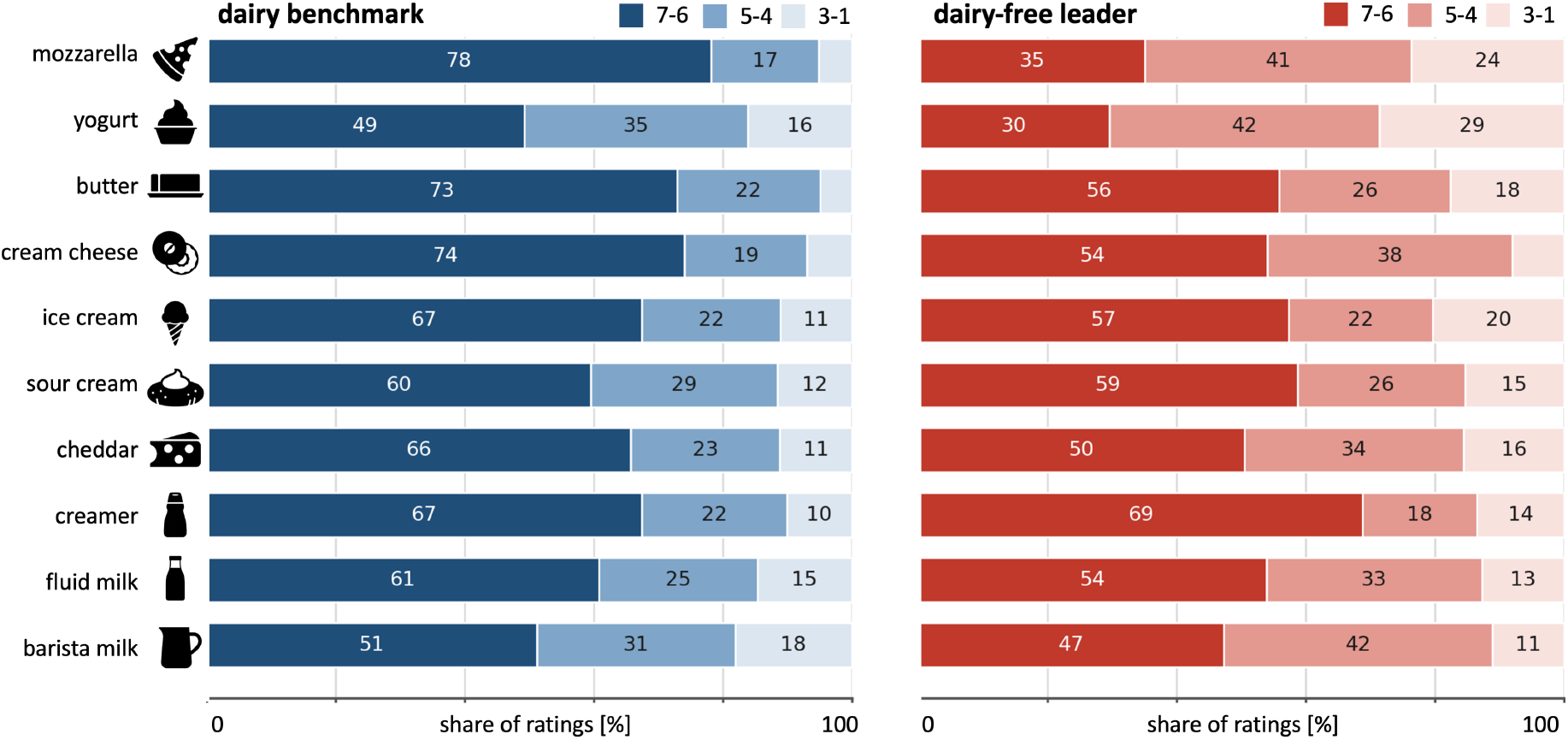
Distribution of overall-liking ratings. Share of participants rating overall liking 6–7 (dark), 4–5 (mid), or 1–3 (light), shown separately for dairy benchmarks (left, blue) and dairy-free leaders (right, red).

### 3.5. Overall liking across age groups

Paired overall-liking differences showed no clear association with age group (*p* = 0.611), and the category-by-age interaction also lacked clear evidence (*p* = 0.431). No within-category age test reached significance after Holm correction. Age subgroups within categories contained 9–38 respondents, which limits precision, and these results do not establish equivalence between age groups (Table S3).

### 3.6. Purchase intent after nutrition-panel reveal

Purchase intent, recorded after participants viewed the nutrition facts panel and ingredient list, separated the categories differently from liking (Table 3, Fig. 5). Six paired purchase-intent deficits survived Holm correction: mozzarella (−1.54), sour cream (−1.44), yogurt (− 1.37), fluid milk (−1.03), cheddar (− 0.74), and butter (− 0.71). Creamer (− 0.06) and ice cream (+0.20) did not differ. Fluid milk met the exploratory liking-equivalence criterion yet had a purchase-intent deficit (−1.03). Because information exposure was not randomized and no prereveal purchase-intent measure was available, the differences cannot be attributed to the nutrition panel or ingredient list. Stated purchase intent also does not measure actual purchases.

**Table 3:** Purchase intent: paired benchmark–leader comparisons. Mean paired difference, dairy-free leader minus dairy benchmark, on the seven-point purchase-intent scale, recorded after participants viewed the nutrition facts panel and ingredient list, with 95% confidence interval, paired *t*-test *p* value, Holm decision across the ten categories, and Cohen’s *d*_*z*_.

| Category | *n* | Mean Δ (95% CI) | *p<sub>t</sub>* | Holm | *d<sub>z</sub>* |
|---|---|---|---|---|---|
| Mozzarella | 114 | −1.54 (−1.92, −1.15) | <0.001 | sig. | −0.74 |
| Sour cream | 104 | −1.44 (−1.88, −1.00) | <0.001 | sig. | −0.63 |
| Yogurt | 105 | −1.37 (−1.78, −0.96) | <0.001 | sig. | −0.65 |
| Fluid milk | 102 | −1.03 (−1.49, −0.57) | <0.001 | sig. | −0.44 |
| Cheddar | 103 | −0.74 (−1.15, −0.33) | <0.001 | sig. | −0.35 |
| Butter | 102 | −0.71 (−1.14, −0.27) | 0.002 | sig. | −0.32 |
| Barista milk | 108 | −0.57 (−1.04, −0.11) | 0.017 | n.s. | −0.23 |
| Cream cheese | 100 | −0.38 (−0.84, +0.08) | 0.103 | n.s. | −0.16 |
| Creamer | 96 | −0.06 (−0.49, +0.36) | 0.770 | n.s. | −0.03 |
| Ice cream | 103 | +0.20 (−0.28, +0.69) | 0.408 | n.s. | +0.08 |

**Figure 5:**
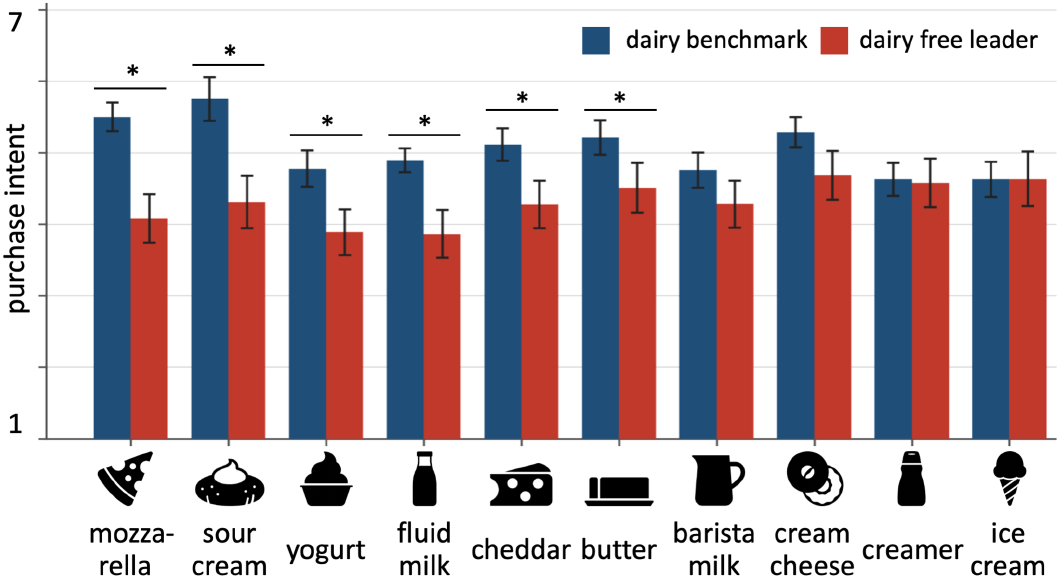
Purchase intent. Paired-subset mean stated purchase intent after participants reviewed the nutrition facts panel and ingredient list (1 = definitely would not buy; 7 = definitely would buy) for the dairy benchmark (blue) and dairy-free leader (red); * paired difference significant after Holm correction. Paired comparisons are reported in Table 3 (*n* = 96–114).

### 3.7. CATA attributes and penalty analysis

The largest category-specific penalties were flavor terms in eight of ten categories (Fig. 6): *off-flavor* in barista milk and cheddar; *artificial*/*chemical* in butter, creamer, and mozzarella; *cardboard*/*stale* in fluid milk; *off-aftertaste* in sour cream; and *bitter* in yogurt; *oily*/*greasy* texture had the largest penalty for ice cream; and *dull* appearance for cream cheese. These estimates rest on small mention counts, and their bootstrap intervals are correspondingly wide; they do not support precise ranking of terms within a category.

**Figure 6:**
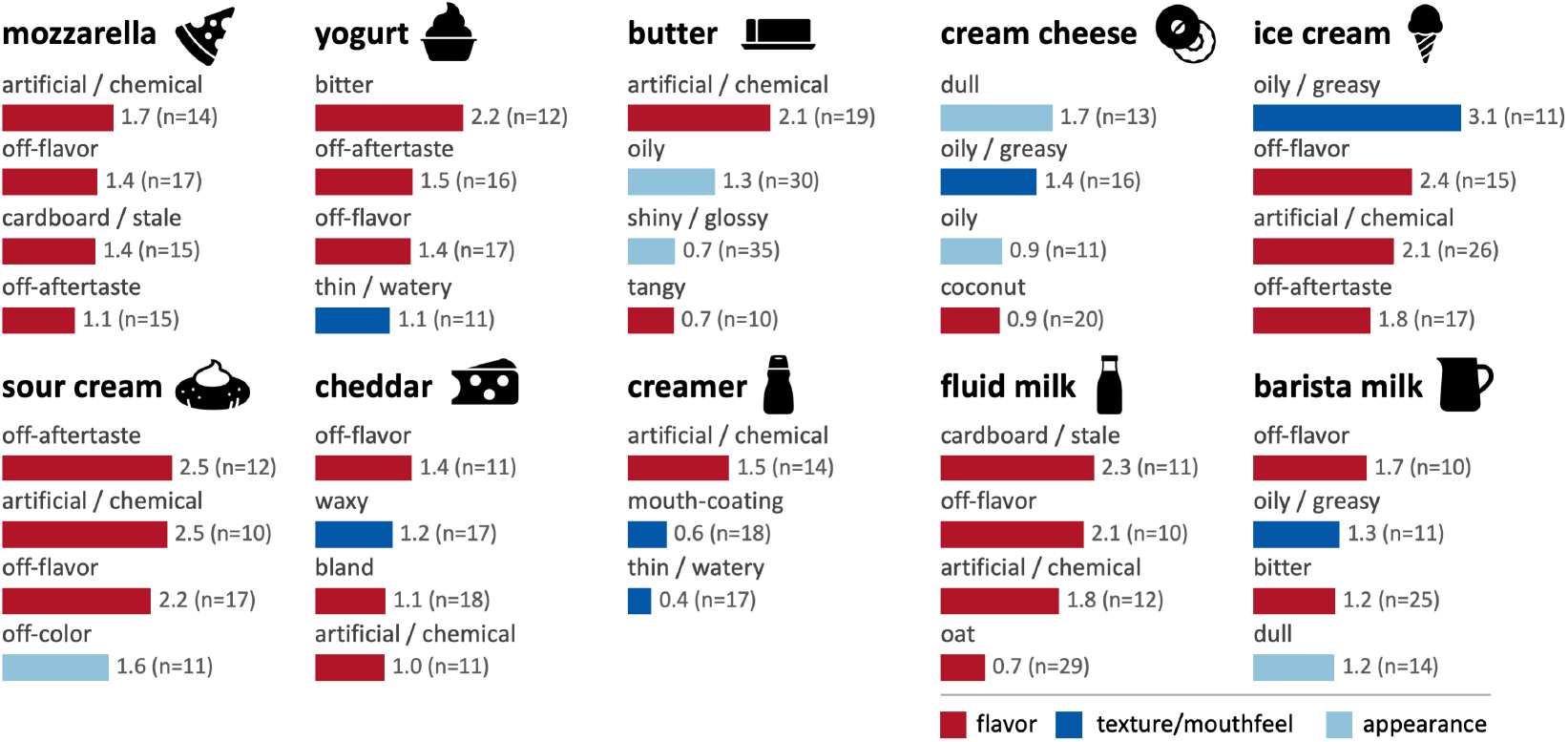
Category-resolved penalty analysis of CATA attributes. Four attributes with the largest penalties for each dairy-free leader among terms selected for at least ten ratings and with penalties of at least 0.3 points. Penalty denotes mean overall liking among participants who did not select the attribute minus the corresponding mean among those who did. Error bars are 95% percentile bootstrap intervals from 10,000 resamples. Bars indicate flavor (red), texture and mouthfeel (blue), or appearance (light blue); labels report mention counts.

In the pooled analysis (Fig. 7), *off-flavor* (penalty 1.85, 95% CI 1.52 to 2.16), *cardboard*/*stale* (1.81, 1.25 to 2.37), and *arti-ficial*/*chemical* (1.80, 1.46 to 2.14) carried the largest penalties.Sensitivity analyses using respondent-clustered bootstrapping and category-adjusted penalties confirmed the robustness of the largest penalties. Respondent clustering did not change which confidence intervals excluded zero, while category adjustment changed this result for only three of 61 terms (Table S4). These descriptive associations generate candidates for follow-up testing, but they do not establish chemical or formulation causes.

**Figure 7:**
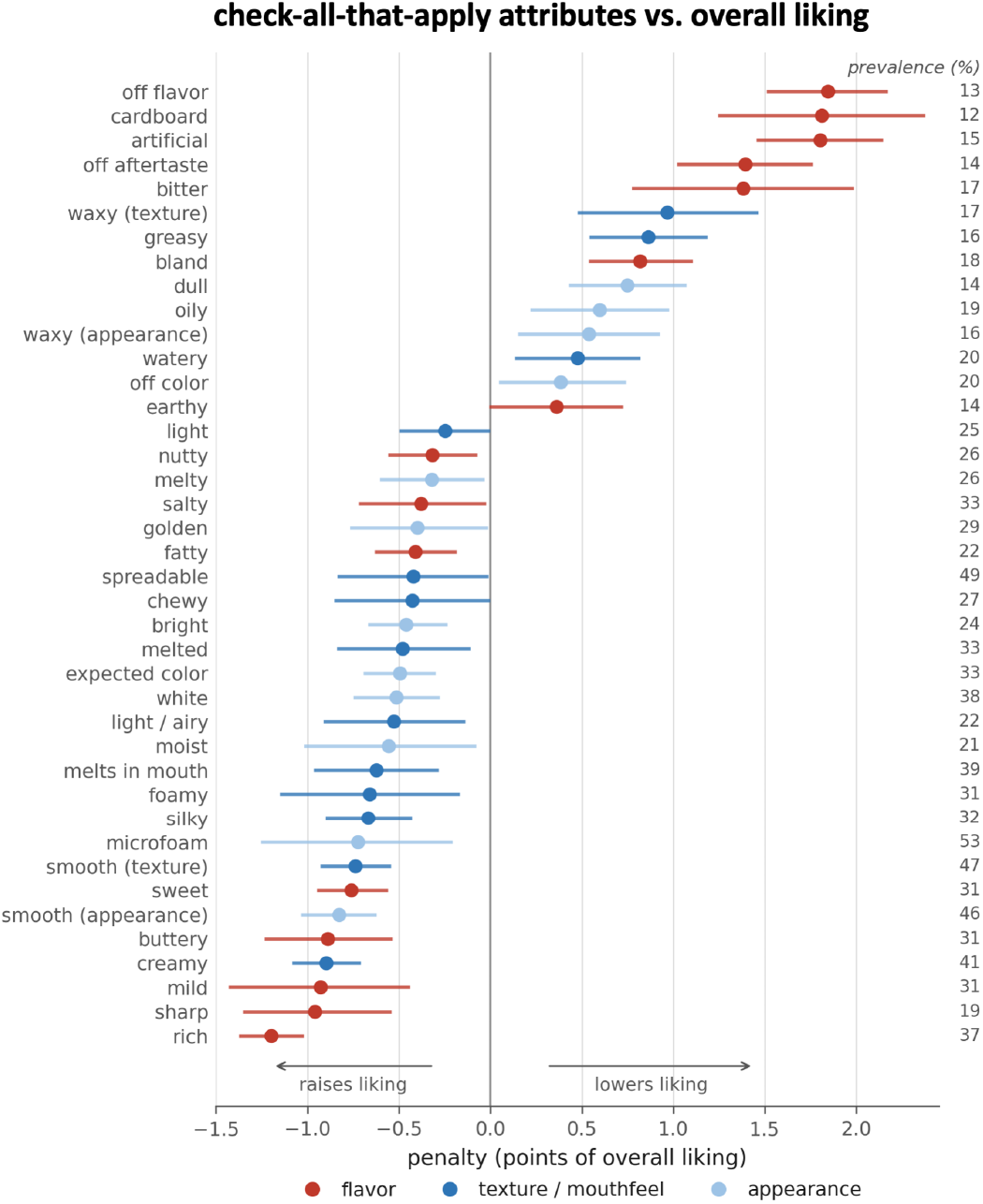
Pooled penalty analysis of CATA attributes. Penalty for each check-all-that-apply attribute pooled across the ten dairy-free leaders. Positive values identify attributes associated with lower liking. Points are ranked by penalty and colored by modality; whiskers are 95% percentile bootstrap intervals from 10,000 resamples. Of the attributes with at least 25 mentions, those whose interval excludes zero are shown; prevalence is printed in the right margin.

### 3.8. Stated price thresholds

The *median bargain*/*expensive-but-still-purchasable* responses were $4.00/$5.00 for sour cream, $4.50/$5.00 for mozzarella, $4.75/$6.00 for creamer, $5.00/$5.50 for cheddar, $5.00/$6.00 for cream cheese, $5.00/$6.50 for fluid milk, $5.50/$6.50 for ice cream, $5.50/$7.00 for barista milk, $5.75/$6.50 for butter, and $7.00/$8.00 for yogurt (*n* = 104 - 418; Fig. 8). Package descriptions differed across categories, so these values should be interpreted only within category.

**Figure 8:**
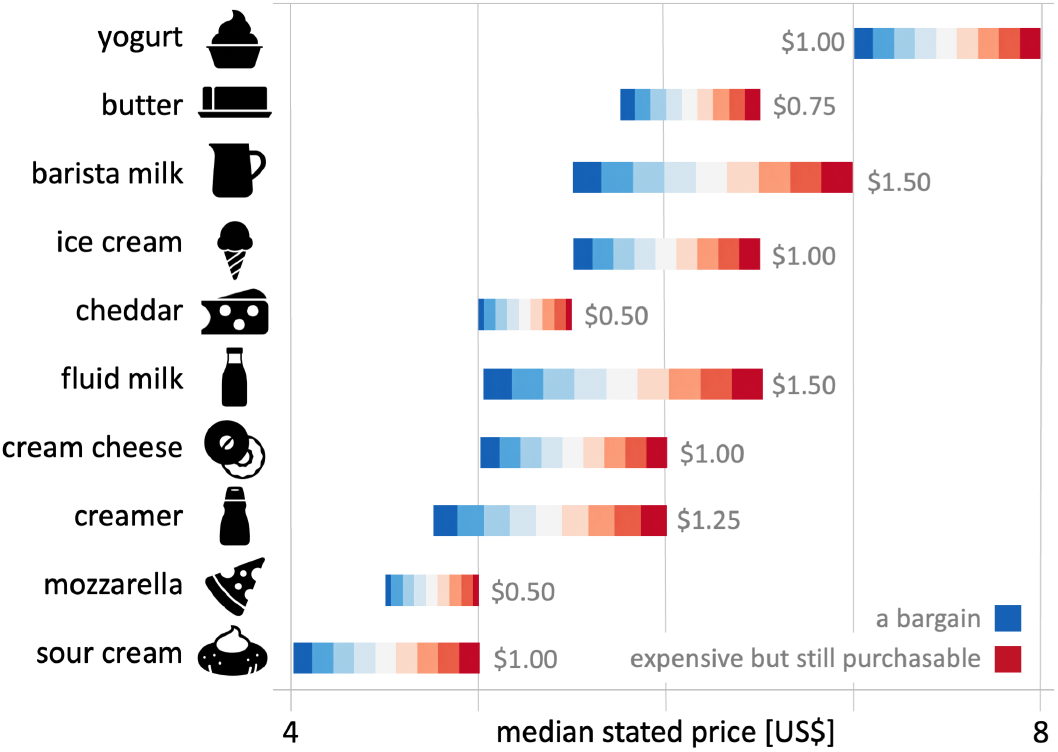
Stated price thresholds. Median prices considered *a bargain* (blue) and *expensive but still purchasable* (red). Package descriptions differed among categories, so comparisons are restricted to the two thresholds within each category.

## 4. Discussion

This study provides a sensory benchmark of selected dairy-free leaders across ten functionally distinct dairy product categories. Unlike most previous sensory studies, which compare a single dairy-free product with its dairy counterpart or focus on a single product category, this work evaluates leading commercial dairy-free products across ten major dairy categories using a common experimental protocol. This unified design enables direct comparisons across applications, identifies category-specific attributes associated with consumer acceptance, and provides quantitative targets for future formulation and instrumental studies.

### Dairy-free leaders perform differently across categories

Mozzarella, yogurt, and butter retained overall-liking deficits after Holm correction, whereas barista milk, creamer, and fluid milk met a chosen ±0.5-point margin in exploratory, unadjusted equivalence tests; none met it after Holm or Bonferroni correction, and the count of categories within the margin depended on the margin. Because that margin was selected for this analysis rather than externally validated, the beverage results support closeness at the stated threshold, not universal or definitive sensory equivalence. The companion meat benchmark likewise found substantial variation among product formats and separation between selected leaders and broader category performance [2]. Both studies therefore describe selected highperforming products, not their markets as a whole.

### Beverages display the smallest liking differences

Coffee can dilute the base beverage and supply dominant roast volatiles, while steaming changes viscosity and foam structure. Protein adsorption, interfacial-film mechanics, continuous-phase viscosity, and liquid drainage are plausible contributors to barista performance [28]. Preparation and serving context can alter the perceptual importance of intrinsic product differences and constrain direct inference from isolated-component measurements to consumer acceptance [23, 11]. Fluid milk also showed a large similarity deficit despite a negligible liking difference, suggesting that acceptance did not require close imitation in this test. These interpretations remain hypotheses because the dataset contains no composition, rheology, foam, or volatile measurements.

### Mozzarella has the largest overall-liking deficit

Its sensory pattern is compatible with the difficulty of combining flow and extensional cohesion in starch-, hydrocolloid-, vegetableoil-, and globular-protein-based matrices [14, 21]. Yogurt and butter present different technical demands involving acid-gel formation, water retention, cultured flavor, fat crystallization, and aroma release [29, 19]. The present consumer data can identify categories and percepts for follow-up, but they cannot demonstrate which formulation mechanism caused a rating difference. Because each leader was selected post hoc as the best-performing product in its category, its component profile need not represent the category. This qualification matters most for mozzarella, where the leader performs comparatively well on texture; the flavor-weighted penalty profile in Fig. 6 should therefore not be read as evidence that texture has been resolved at the category level.

### CATA penalties should be treated as screening associations

They can help identify attributes associated with consumer liking and disliking, but not as evidence of causality. Flavor-related terms had the largest descriptive penalties in most categories, but raw-material volatiles, oxidation, processing, ingredient interactions, and oral release kinetics could all contribute [15, 26]. The small category-specific mention counts, wide bootstrap intervals, lack of adjustment for category, and possible repeated participation prevent causal or ingredient-level attribution. Related consumer studies like-wise identify sensory improvement targets as formulation-and context-specific, supporting attribute penalties as screening tools rather than general rules for product reformulation [10, 25, 24].

### Purchase intent incorporates factors beyond taste

It was measured only after participants viewed the nutrition panel and ingredient list. Six category deficits survived Holm correction, including fluid milk and yogurt. The gap cannot be assigned to the reveal because the study lacks randomized information exposure and a pre-reveal measure. The result shows that stated purchase intent at that stage can diverge from blinded liking; it does not identify whether nutrition information, ingredients, expected price, prior experience, or another factor explains the divergence [20]. Stated intent also does not measure actual purchase behavior.

### Limitations

Three limitations qualify our findings: First, NECTAR selected category leaders post hoc from multiple candidates, but the available data do not permit correction for selection effects. Component ratings and CATA analyses remain exploratory, and none of the category-specific equivalence findings survives multiplicity correction. The primary pooled boot-strap does not account for category or repeated participation, including 170 respondents who contributed pairs in multiple categories; yet, respondent-clustered and category-adjusted sensitivity analyses support the robustness of the largest penalties. Second, our findings reflect complete products under specific serving conditions and a panel of regular category consumers rather than a representative population. Third, the study lacks compositional, microstructural, chemical, and mechanical measurements to explain the observed sensory differences. Products also differ in water and nutrient content; similar sensory ratings therefore do not establish nutritional equivalence. Controlled sensory and instrumental studies can test the proposed mechanisms and connect product composition and structure to consumer perception.

## 5. Conclusions

This study establishes the first open-source sensory benchmark that compares selected *dairy-free leaders* with *dairy benchmarks* across ten major food categories under blinded evaluation. Rather than asking whether dairy-free products perform well in general, it quantifies category-specific sensory gaps and identifies sensory attributes associated with lower liking. The resulting benchmark offers a standardized reference that researchers and product developers can use to evaluate commercial products and emerging dairy-free formulations. Sensory performance varied substantially across product categories. The selected dairy-free leaders closely approached conventional dairy in several beverage applications. Barista milk, creamer, and fluid milk met the chosen exploratory ±0.5-point equivalence criterion without multiplicity adjustment, although none met it after Holm or Bonferroni correction, while cheddar, sour cream, and ice cream differed by less than 0.3 points in mean overall liking. By contrast, mozzarella showed the largest sensory gap of −1.46 points on the 7-point overall-liking scale, and yogurt and butter also remained significantly different after Holm correction. Descriptive CATA analyses associated off-flavor, artificial or chemical notes, bitter taste, and lingering aftertaste with lower liking, and these associations suggest candidates for follow-up tests. Together, these findings indicate that the selected dairy-free beverages approached the liking of their dairy benchmarks, while structured products such as mozzarella, yogurt, and butter still offer substantial opportunities for improvement. Priorities for controlled sensory and instrumental studies include foam stability in barista milk, melt and stretch in mozzarella, and gel texture in cultured products. Beyond the category-specific results, the benchmark dataset, available through NECTAR, provides a quantitative foundation to connect formulation, instrumental characterization, and consumer perception. It also establishes a common reference that can guide the development of dairy-free products with sensory quality that rivals conventional dairy.

## CRediT authorship contribution statement

Aeneas O. Koosis: Formal analysis, visualization, writing - original draft, writing - review and editing. Ellen Kuhl: Conceptualization, methodology, supervision, funding acquisition, writing - original draft, writing - review and editing. Caroline Cotto: Conceptualization, investigation, resources, data curation, project administration, writing - review and editing.

## Ethics statement

Palate Insights collected the sensory and demographic data under informed consent procedures and applicable privacy safe-guards as part of the NECTAR Taste of the Industry 2026 analysis. This study represents a secondary analysis of the resulting fully de-identified dataset. Stanford University determined that this research does not involve human subjects, as defined in 45 CFR 46.102(e).

## Informed consent

Informed consent was obtained from all subjects involved in the study.

## Data availability

NECTAR provides the data at https://www.nectar.org/sensory-research/2026-taste-of-the-industry after registration through its research platform. NECTAR manages access and any wider public release. The analysis code is available on the Living Matter Lab GitHub at https://github.com/LivingMatterLab.

## Acknowledgments

The sensory survey was conducted by Palate Insights. The authors thank Alex Weissman, Han Gu, and Tom Conger from Palate Insights and Max Elder from Food System Innovations for their guidance and support.

## Declaration of generative AI and AI-assisted technologies in the manuscript preparation process

During the preparation of this work, the authors used AI-assisted tools to support manuscript organization, drafting, editing, and code development. After using these tools, the authors reviewed and edited the content as needed and take full responsibility for the content of the published article.

## Supplementary material

**Table S1:** Equivalence tests for overall liking at the *±*0.5-point margin. Mean paired difference, dairy-free leader minus dairy benchmark, with the 90% confidence interval used for the TOST decision. Holm and Bonferroni adjustments span the ten categories. A category lies within the margin when its *p*_TOST_ *<* 0.05. Categories are ordered by mean difference.

| Category | *n* | Mean Δ | 90% CI | *p*<sub>TOST</sub> |  |  | Within margin |  |
|---|---|---|---|---|---|---|---|---|
|  |  |  |  | Unadjusted | Holm | Bonferroni | Unadjusted | Holm |
| Mozzarella | 114 | −1.46 | (−1.71, −1.20) | 1.000 | 1.000 | 1.000 | no | no |
| Yogurt | 105 | −0.68 | (−1.03, −0.32) | 0.793 | 1.000 | 1.000 | no | no |
| Butter | 102 | −0.54 | (−0.83, −0.25) | 0.589 | 1.000 | 1.000 | no | no |
| Cream cheese | 100 | −0.42 | (−0.71, −0.13) | 0.326 | 1.000 | 1.000 | no | no |
| Ice cream | 103 | −0.28 | (−0.68, +0.11) | 0.181 | 0.906 | 1.000 | no | no |
| Sour cream | 104 | −0.26 | (−0.60, +0.08) | 0.122 | 0.805 | 1.000 | no | no |
| Cheddar | 103 | −0.25 | (−0.59, +0.09) | 0.115 | 0.805 | 1.000 | no | no |
| Creamer | 96 | −0.06 | (−0.37, +0.25) | 0.011 | 0.106 | 0.106 | yes | no |
| Fluid milk | 102 | +0.02 | (−0.35, +0.39) | 0.017 | 0.152 | 0.169 | yes | no |
| Barista milk | 108 | +0.17 | (−0.14, +0.48) | 0.038 | 0.306 | 0.382 | yes | no |

**Table S2:** Sensitivity of the equivalence results to the margin. Unadjusted TOST *p* values for overall liking at seven margins on the seven-point scale. Bold values meet the unadjusted criterion (*p<* 0.05); an asterisk marks values that also meet it after Holm adjustment across the ten categories. The final rows count the categories that meet each criterion.

| Category | ±0.25 | ±0.3 | ±0.4 | ±0.5 | ±0.6 | ±0.75 | ±1.0 |
|---|---|---|---|---|---|---|---|
| Mozzarella | 1.000 | 1.000 | 1.000 | 1.000 | 1.000 | 1.000 | 0.998 |
| Yogurt | 0.975 | 0.958 | 0.899 | 0.793 | 0.638 | 0.366 | 0.068 |
| Butter | 0.951 | 0.915 | 0.788 | 0.589 | 0.363 | 0.113 | **0.005**\* |
| Cream cheese | 0.830 | 0.750 | 0.545 | 0.326 | 0.156 | **0.033** | <**0.001**\* |
| Ice cream | 0.552 | 0.469 | 0.310 | 0.181 | 0.093 | **0.026** | **0.002**\* |
| Sour cream | 0.519 | 0.422 | 0.248 | 0.122 | 0.050 | **0.009** | <**0.001**\* |
| Cheddar | 0.505 | 0.408 | 0.237 | 0.115 | **0.047** | **0.008** | <**0.001**\* |
| Creamer | 0.159 | 0.103 | **0.037** | **0.011** | **0.002**\* | <**0.001**\* | <**0.001**\* |
| Fluid milk | 0.152 | 0.106 | **0.046** | **0.017** | **0.005**\* | <**0.001**\* | <**0.001**\* |
| Barista milk | 0.328 | 0.238 | 0.107 | **0.038** | **0.011** | **0.001**\* | <**0.001**\* |
| Categories within margin, unadjusted | 0 | 0 | 2 | 3 | 4 | 7 | 8 |
| Categories within margin, Holm | 0 | 0 | 0 | 0 | 2 | 3 | 8 |
| Categories within margin, Bonferroni | 0 | 0 | 0 | 0 | 1 | 3 | 8 |

**Table S3:**
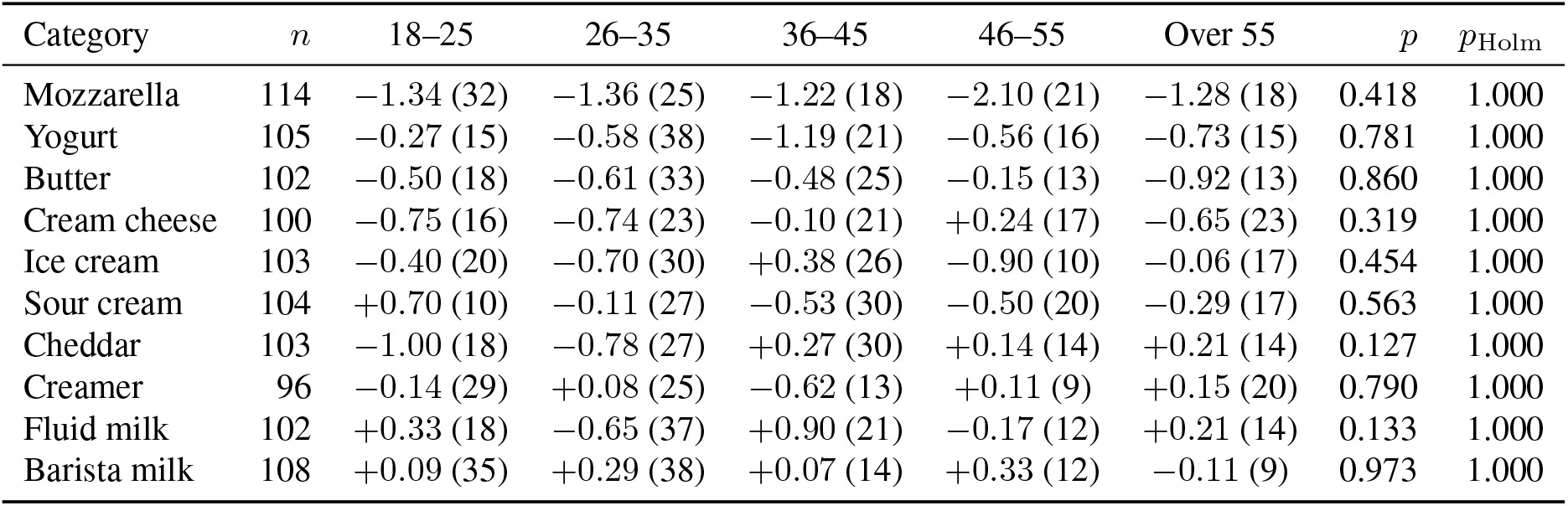
Overall-liking differences by age group. Mean paired difference, dairy-free leader minus dairy benchmark, with the number of respondents in parentheses. The analysis includes 1,037 paired observations from 725 respondents; 170 respondents contributed pairs in more than one category. A linear model with category and age group as factors and respondent-clustered standard errors showed no clear age effect (*F*_4,724_ = 0.67, *p* = 0.611) and no clear category-by-age interaction (*F*_36,724_ = 1.02, *p* = 0.431). Within-category *p* values come from one-way ANOVA across the five age groups, with Holm adjustment across the ten categories. Subgroups contain 9–38 respondents, which limits precision, and these results do not establish equivalence between age groups.

**Table S4:** Pooled CATA penalties with respondent-clustered and category-adjusted estimates. Penalty denotes mean overall liking among leader ratings without the term minus the mean among ratings with it; positive values mark terms associated with lower liking. The original analysis resamples ratings within the selected and unselected groups. The respondent-clustered bootstrap resamples respondents, so it keeps all ratings from a respondent together. The category-adjusted penalty averages the within-category penalties, weighted by *n*_sel_*n*_unsel_*/*(*n*_sel_ + *n*_unsel_), so differences in liking between categories do not contribute; its interval comes from the same respondent-clustered bootstrap. Each bootstrap uses 10,000 resamples. The final column gives the range of within-category penalties. Intervals exclude zero for 40, 40,and 39 of 61 terms in the original, clustered, and category-adjusted analyses. Terms are ordered by category-adjusted penalty.

|  |  |  |  | Pooled penalty |  |  | Category-adjusted |  |  |
|---|---|---|---|---|---|---|---|---|---|
| Term | Modality | Mentions | Categories | Penalty | Original 95% CI | Clustered 95% CI | Penalty | Clustered 95% CI | Category range |
| Artificial / Chemical | flavor | 106 | 7 | +1.80 | (+1.46, +2.14) | (+1.45, +2.14) | +1.85 | (+1.50, +2.20) | +1.04 to +2.47 |
| Off-flavor | flavor | 97 | 7 | +1.85 | (+1.52, +2.16) | (+1.53, +2.16) | +1.82 | (+1.50, +2.14) | +1.43 to +2.38 |
| Cardboard / Stale | flavor | 26 | 2 | +1.81 | (+1.25, +2.37) | (+1.24, +2.36) | +1.79 | (+1.20, +2.35) | +1.40 to +2.30 |
| Bitter | flavor | 37 | 2 | +1.38 | (+0.78, +1.98) | (+0.77, +1.99) | +1.58 | (+1.00, +2.13) | +1.23 to +2.22 |
| Off-aftertaste | flavor | 87 | 6 | +1.39 | (+1.03, +1.75) | (+1.04, +1.76) | +1.38 | (+1.03, +1.75) | +0.70 to +2.55 |
| Waxy | texture | 37 | 2 | +0.97 | (+0.48, +1.46) | (+0.47, +1.45) | +0.96 | (+0.47, +1.44) | +0.78 to +1.16 |
| Oily / Greasy | texture | 136 | 8 | +0.86 | (+0.55, +1.18) | (+0.55, +1.18) | +0.93 | (+0.60, +1.25) | +0.16 to +3.11 |
| Bland | flavor | 130 | 7 | +0.82 | (+0.55, +1.10) | (+0.54, +1.10) | +0.77 | (+0.50, +1.05) | +0.26 to +1.08 |
| Dull | appearance | 120 | 8 | +0.75 | (+0.44, +1.06) | (+0.44, +1.06) | +0.72 | (+0.41, +1.04) | −0.39 to +1.68 |
| Oily | appearance | 100 | 5 | +0.60 | (+0.23, +0.97) | (+0.23, +0.97) | +0.63 | (+0.27, +1.02) | +0.17 to +1.31 |
| Thin / Watery | texture | 101 | 5 | +0.48 | (+0.14, +0.81) | (+0.14, +0.82) | +0.57 | (+0.23, +0.90) | +0.18 to +1.12 |
| Waxy | appearance | 85 | 5 | +0.54 | (+0.16, +0.92) | (+0.15, +0.91) | +0.48 | (+0.09, +0.86) | +0.06 to +0.84 |
| Gummy / Gooey | texture | 48 | 3 | +0.43 | (−0.10, +0.97) | (−0.10, +0.98) | +0.45 | (−0.06, +0.98) | −0.20 to +1.09 |
| Earthy / Grassy | flavor | 88 | 6 | +0.36 | (+0.00, +0.71) | (+0.01, +0.73) | +0.38 | (+0.03, +0.74) | +0.00 to +1.06 |
| Off-color | appearance | 127 | 6 | +0.38 | (+0.06, +0.73) | (+0.04, +0.72) | +0.37 | (+0.02, +0.71) | −0.42 to +1.60 |
| Partially Unmelted | texture | 47 | 2 | +0.14 | (−0.26, +0.55) | (−0.27, +0.55) | +0.26 | (−0.16, +0.67) | −0.01 to +0.43 |
| Partially Unmelted | appearance | 47 | 2 | +0.03 | (−0.41, +0.49) | (−0.42, +0.49) | +0.24 | (−0.23, +0.74) | −0.17 to +0.40 |
| Greyish | appearance | 71 | 2 | +0.16 | (−0.30, +0.64) | (−0.31, +0.65) | +0.19 | (−0.33, +0.71) | −0.25 to +0.44 |
| Uniform | appearance | 39 | 1 | +0.13 | (−0.47, +0.73) | (−0.45, +0.75) | +0.13 | (−0.45, +0.75) | +0.13 to +0.13 |
| Orange | appearance | 45 | 1 | +0.13 | (−0.44, +0.69) | (−0.46, +0.69) | +0.13 | (−0.46, +0.69) | +0.13 to +0.13 |
| Mouth-coating | texture | 201 | 9 | +0.14 | (−0.12, +0.40) | (−0.14, +0.42) | +0.12 | (−0.16, +0.40) | −0.37 to +0.58 |
| Yellowish | appearance | 43 | 2 | +0.12 | (−0.37, +0.60) | (−0.37, +0.61) | +0.09 | (−0.40, +0.59) | −0.09 to +0.22 |
| Coconut | flavor | 102 | 5 | +0.13 | (−0.21, +0.48) | (−0.23, +0.48) | +0.06 | (−0.30, +0.41) | −0.46 to +0.89 |
| Dense / Heavy | texture | 97 | 6 | +0.06 | (−0.29, +0.43) | (−0.27, +0.42) | +0.06 | (−0.27, +0.40) | −0.35 to +0.67 |
| Shiny / Glossy | appearance | 269 | 10 | −0.00 | (−0.22, +0.22) | (−0.22, +0.23) | +0.01 | (−0.20, +0.24) | −0.63 to +0.71 |
| Sour | flavor | 67 | 2 | −0.11 | (−0.59, +0.39) | (−0.61, +0.40) | +0.01 | (−0.50, +0.50) | −0.63 to +0.79 |
| Springy | texture | 33 | 2 | +0.06 | (−0.43, +0.58) | (−0.45, +0.59) | −0.03 | (−0.56, +0.51) | −0.26 to +0.45 |
| Thick | appearance | 112 | 3 | +0.07 | (−0.30, +0.45) | (−0.30, +0.45) | −0.08 | (−0.46, +0.29) | −0.13 to −0.06 |
| Thick | texture | 149 | 6 | −0.03 | (−0.33, +0.28) | (−0.32, +0.28) | −0.14 | (−0.44, +0.17) | −0.55 to +0.21 |
| Browning | appearance | 28 | 1 | −0.15 | (−0.80, +0.49) | (−0.80, +0.51) | −0.15 | (−0.80, +0.51) | −0.15 to −0.15 |
| Whitens Coffee | appearance | 73 | 2 | −0.19 | (−0.58, +0.21) | (−0.58, +0.21) | −0.17 | (−0.57, +0.23) | −0.28 to −0.07 |
| Yellow | appearance | 103 | 2 | −0.23 | (−0.65, +0.19) | (−0.65, +0.20) | −0.23 | (−0.72, +0.26) | −0.43 to −0.02 |
| Light | texture | 185 | 7 | −0.25 | (−0.49, −0.01) | (−0.48, −0.01) | −0.25 | (−0.49, −0.01) | −1.32 to +0.45 |
| Salty | flavor | 105 | 3 | −0.38 | (−0.71, −0.04) | (−0.72, −0.03) | −0.27 | (−0.65, +0.12) | −0.55 to −0.03 |
| Nutty | flavor | 216 | 8 | −0.32 | (−0.55, −0.08) | (−0.55, −0.08) | −0.32 | (−0.56, −0.07) | −0.80 to +0.07 |
| Oat | flavor | 114 | 5 | −0.28 | (−0.61, +0.05) | (−0.60, +0.05) | −0.32 | (−0.64, +0.02) | −1.01 to +0.67 |
| Tangy | flavor | 126 | 6 | −0.27 | (−0.54, +0.02) | (−0.56, +0.04) | −0.32 | (−0.62, −0.01) | −0.90 to +0.69 |
| Soft | texture | 90 | 2 | −0.34 | (−0.80, +0.12) | (−0.79, +0.13) | −0.33 | (−0.79, +0.13) | −0.42 to −0.23 |
| Bright | appearance | 222 | 9 | −0.46 | (−0.66, −0.25) | (−0.67, −0.24) | −0.38 | (−0.59, −0.16) | −0.83 to +0.10 |
| Golden | appearance | 91 | 3 | −0.40 | (−0.76, −0.02) | (−0.78, −0.01) | −0.38 | (−0.76, +0.01) | −0.62 to −0.08 |
| Fatty | flavor | 228 | 10 | −0.41 | (−0.62, −0.19) | (−0.63, −0.19) | −0.40 | (−0.62, −0.18) | −1.38 to +0.62 |
| White | appearance | 275 | 7 | −0.51 | (−0.74, −0.29) | (−0.74, −0.28) | −0.42 | (−0.65, −0.18) | −0.89 to −0.05 |
| Spreadable | texture | 99 | 2 | −0.42 | (−0.83, −0.02) | (−0.84, −0.00) | −0.43 | (−0.85, −0.01) | −0.44 to −0.42 |
| Melty | appearance | 162 | 6 | −0.32 | (−0.60, −0.04) | (−0.58, −0.04) | −0.44 | (−0.70, −0.17) | −1.20 to −0.07 |
| Expected Color | appearance | 347 | 10 | −0.49 | (−0.68, −0.31) | (−0.68, −0.31) | −0.45 | (−0.63, −0.26) | −1.08 to +0.06 |
| Melted | texture | 105 | 3 | −0.48 | (−0.83, −0.12) | (−0.83, −0.12) | −0.50 | (−0.85, −0.14) | −1.11 to −0.10 |
| Light / Airy | texture | 68 | 3 | −0.53 | (−0.90, −0.15) | (−0.90, −0.16) | −0.53 | (−0.89, −0.19) | −0.98 to −0.38 |
| Moist | appearance | 42 | 2 | −0.56 | (−1.01, −0.09) | (−1.03, −0.06) | −0.55 | (−1.03, −0.06) | −0.78 to −0.32 |
| Chewy | texture | 58 | 2 | −0.43 | (−0.84, −0.01) | (−0.84, −0.01) | −0.58 | (−1.00, −0.17) | −0.63 to −0.50 |
| Melts-in-mouth | texture | 120 | 3 | −0.62 | (−0.96, −0.29) | (−0.96, −0.28) | −0.64 | (−0.99, −0.29) | −0.74 to −0.57 |
| Silky | texture | 229 | 7 | −0.67 | (−0.89, −0.44) | (−0.90, −0.43) | −0.66 | (−0.89, −0.41) | −0.98 to −0.14 |
| Foamy | texture | 34 | 1 | −0.66 | (−1.14, −0.18) | (−1.16, −0.15) | −0.66 | (−1.16, −0.15) | −0.66 to −0.66 |
| Smooth | texture | 485 | 10 | −0.74 | (−0.92, −0.55) | (−0.93, −0.54) | −0.70 | (−0.90, −0.50) | −1.26 to −0.22 |
| Microfoam | appearance | 57 | 1 | −0.72 | (−1.25, −0.22) | (−1.24, −0.20) | −0.72 | (−1.24, −0.20) | −0.72 to −0.72 |
| Sweet | flavor | 326 | 10 | −0.76 | (−0.94, −0.57) | (−0.95, −0.56) | −0.75 | (−0.96, −0.55) | −1.13 to −0.37 |
| Smooth | appearance | 432 | 9 | −0.83 | (−1.03, −0.63) | (−1.03, −0.63) | −0.79 | (−1.00, −0.59) | −1.18 to −0.37 |
| Buttery | flavor | 97 | 3 | −0.89 | (−1.23, −0.54) | (−1.23, −0.54) | −0.89 | (−1.23, −0.55) | −1.16 to −0.51 |
| Sharp | flavor | 42 | 2 | −0.96 | (−1.34, −0.55) | (−1.36, −0.54) | −0.91 | (−1.32, −0.48) | −1.34 to −0.58 |
| Creamy | texture | 428 | 10 | −0.90 | (−1.07, −0.72) | (−1.07, −0.72) | −0.92 | (−1.10, −0.75) | −1.11 to −0.60 |
| Mild | flavor | 32 | 1 | −0.93 | (−1.42, −0.45) | (−1.42, −0.43) | −0.93 | (−1.42, −0.43) | −0.93 to −0.93 |
| Rich | flavor | 386 | 10 | −1.20 | (−1.37, −1.03) | (−1.36, −1.03) | −1.15 | (−1.32, −0.99) | −1.80 to −0.84 |

## References

[1] Alsado, C., Chen, L., Wismer, W., 2025. Are plant-based milks unique products or dairy milk substitutes? A study of consumer perceptions, uses, and consumption motivations of plant-based and dairy milks. Food Quality and Preference 134, 105681. doi:10.1016/j.foodqual.2025.105681.foods15122112.

[2] van den Bedem, S.D., Kuhl, E., Cotto, C., 2026. Open-source bench-marking of plant-based and animal meats. Foods 15, 2112. doi:10.3390/

[3] Bertini, A., Cáceres-Ozimica, H., Valenzuela, R., Durán-Agöero, S., 2026. Global and regional inequalities in dairy recommendations: a natural language processing analysis of food-based dietary guidelines across income groups. The Lancet Regional Health – Americas 62, 101550. doi:10.1016/j.lana.2026.101550.

[4] Catanzaro, R., Sciuto, M., Marotta, F., 2021. Lactose intolerance—old and new knowledge on pathophysiological mechanisms, diagnosis, and treatment. SN Comprehensive Clinical Medicine 3, 499–509. doi:10.1007/s42399-021-00792-9.

[5] Clark, M.A., Domingo, N.G.G., Colgan, K., Thakrar, S.K., Tilman, D., Lynch, J., Azevedo, I.L., Hill, J.D., 2020. Global food system emissions could preclude achieving the 1.5↔ and 2↔C climate change targets. Science 370, 705–708. doi:10.1126/science.aba7357.

[6] Crippa, M., Solazzo, E., Guizzardi, D., Monforti-Ferrario, F., Tubiello, F.N., Leip, A., 2021. Food systems are responsible for a third of global anthropogenic GHG emissions. Nature Food 2, 198–209. doi:10.1038/s43016-021-00225-9.

[7] Good Food Institute, 2026. U.S. retail market insights for the plant-based industry. The Good Food Institute. 2025 SPINS retail sales data. Available at https://gfi.org/marketresearch/.

[8] Jaeger, S.R., Dupas de Matos, A., Frempomaa Oduro, A., Hort, J., 2024. Sensory characteristics of plant-based milk alternatives: product characterisation by consumers and drivers of liking. Food Research International 180, 114093. doi:10.1016/j.foodres.2024.114093.

[9] Khanpit, V., Viswanathan, S., Hinrichsen, O., 2024. Environmental impact of animal milk vs plant-based milk: critical review. Journal of Cleaner Production 449, 141703. doi:10.1016/j.jclepro.2024.141703.

[10] Koosis, A.O., Gardner, C.D., Kuhl, E., 2026. The burger battle: The future is blended. bioRxiv URL: https://www.biorxiv.org/content/10.64898/2026.06.17.732995v1, doi:10.64898/2026.06.17.732995. preprint, June 17, 2026.

[11] Koosis, A.O., Kuhl, E., 2026. Texture profile analysis and consumer sensory evaluation of plant-based and conventional breaded shrimp. bioRxiv doi:10.64898/2026.07.17.739214. preprint, posted 20 July 2026.

[12] Kuhl, E., 2025. AI for food: accelerating and democratizing discovery and innovation. npj Science of Food 9, 82. doi:10.1038/s41538-025-00441-8.s11947-024-03446-5.

[13] Li, X., Zhou, S., Chen, H., Zhang, R., Wang, L., 2025. Pomelo fiberstabilized oil-in-water emulsion gels: fat mimetic in plant-based ice cream. Food and Bioprocess Technology 18, 422–432. doi:10.1007/

[14] Mattice, K.D., Marangoni, A.G., 2020. Physical properties of plant-based cheese products produced with zein. Food Hydrocolloids 105, 105746. doi:10.1016/j.foodhyd.2020.105746.

[15] McClements, D.J., Grossmann, L., 2021. The science of plant-based foods: constructing next-generation meat, fish, milk, and egg analogs. Comprehensive Reviews in Food Science and Food Safety 20, 4049–4100. doi:10.1111/1541-4337.12771.

[16] National Health Service, 2023. Dairy and alternatives in your https://www.nhs.uk/live-well/eat-well/food-types/milk-and-dairy-nutrition/. Last reviewed 11 December 2023.

[17] NECTAR, 2026. Taste of the Industry 2026: A Sensory Analysis of Dairy-Free Products. Technical Report. Food System Innovations. Claremont, CA, USA. Creative Commons Attribution-NonCommercial 4.0 International License.

[18] Poore, J., Nemecek, T., 2018. Reducing food’s environmental impacts through producers and consumers. Science 360, 987–992. doi:10.1126/science.aaq0216.

[19] Rønholt, S., Kirkensgaard, J.J.K., Pedersen, T.B., Mortensen, K., Knudsen, J.C., 2012. Polymorphism, microstructure and rheology of butter: effects of cream heat treatment. Food Chemistry 135, 1730–1739. doi:10.1016/j.foodchem.2012.05.087.

[20] Rouf, A., Babu, A., Siddikee, J.A., Islam, R., Afrin, S., Hasan, M., Jaman, S.S., 2025. The careful consumer: effects of altruistic and egoistic motivation on the purchase intention of green products. Quality & Quantity 59, 3353–3375. doi:10.1007/s11135-025-02123-9.

[21] Rune, C.J.B., Clausen, M.P., Giacalone, D., 2026. Sensory evaluation of plant-based cheese: a systematic review with a focus on texture and mouthfeel. Critical Reviews in Food Science and Nutrition 66, 754–779. doi:10.1080/10408398.2025.2531220.

[22] Springmann, M., Clark, M., Mason-D’Croz, D., Wiebe, K., Bodirsky, B.L., Lassaletta, L., et al., 2018. Options for keeping the food system within environmental limits. Nature 562, 519–525. doi:10.1038/s41586-018-0594-0.

[23] St. Pierre, S.R., Darwin, E.C., Adil, D., Aviles, M.C., Date, A., Dunne, R.A., Lall, Y., Parra Vallecillo, M., Perez Medina, V.A., Linka, K., Levenston, M.E., Kuhl, E., 2024. The mechanical and sensory signature of plant-based and animal meat. npj Science of Food 8, 94. doi:10.1038/s41538-024-00330-6.

[24] St. Pierre, S.R., Koosis, A., Zhang, N., Kuhl, E., 2026. The meatball matchup: plant vs. animal proteins on campus. Food Research International 240, 119631. doi:10.1016/j.foodres.2026.119631.

[25] Tac, V., Koosis, A.O., Kuhl, E., 2026. Texture independently drives liking in AI-generated alternative protein burgers. Foods 15, 2026. doi:10.3390/foods15112026.

[26] Tian, H., Yan, X., Fu, L., Hang, S., Yu, H., Chen, C., Ge, C., 2026. Flavor formation mechanisms and modulation strategies in plant-based cheese analogs: from raw material selection to technological innovations. Comprehensive Reviews in Food Science and Food Safety 25, e70537. doi:10.1111/1541-4337.70537.

[27] Willett, W., Rockström, J., Loken, B., Springmann, M., Lang, T., Vermeulen, S., et al., 2019. Food in the Anthropocene: the EAT–Lancet Commission on healthy diets from sustainable food systems. The Lancet 393, 447–492. doi:10.1016/S0140-6736(18)31788-4.

[28] Wöest, S., Buczkowski, J., Jones, N.C., Hoffmann, S.V., Fischer, P., Wooster, T.J., 2025. Plant vs dairy protein stabilised cappuccino foams: how protein and hydrocolloid conformational changes affect foam stability. Food Hydrocolloids 169, 111621. doi:10.1016/j.foodhyd.2025.111621.

[29] Yin, X., Li, J., Zhu, L., Zhang, H., 2024. Advances in the formation mechanism of set-type plant-based yogurt gel: a review. Critical Reviews in Food Science and Nutrition 64, 9412–9431. doi:10.1080/10408398.2023.2212764.

